# Cell context-specific Synthetic lethality Prediction and Mechanism Analysis

**DOI:** 10.1101/2023.09.13.557545

**Authors:** Yucui Xing, Mengchen Pu, Kaiyang Cheng, Kai Tian, Lanying Wei, Weisheng Zheng, Gongxin Peng, Jielong Zhou, Yingsheng Zhang

## Abstract

Synthetic lethality (SL) holds significant promise as a targeted cancer therapy by selectively eliminating tumor cells while sparing normal cells. However, the discovery of SL gene pairs has encountered tremendous challenges, including high costs and low efficiency of experimental methods. Current computational approaches only provide limited insights because of overlooking the crucial aspects of cellular context dependency and mechanistic understanding of SL pairs. To overcome these challenges, we have developed SLWise, a deep-learning model capable of predicting SL interactions in diverse cellular backgrounds. We evaluated SLWise using a real world ground truth standard. The evaluation demonstrated that SLWise outperformed benchmark models in SL prediction. Additionally, we proposed a novel analysis scheme called SLAD-CE (**S**ynthetic **L**ethal **A**ssociated Gene **D**etection and **C**ell Damage **E**valuation) for the identification of abnormal essential genes induced by SL gene pairs and tracking the extent of cell damage. Leveraging the cell-line-specific input feature L1000 and employing Gene Set Enrichment Analysis (GSEA), SLAD-CE provides valuable insights into the underlying mechanisms of SLWise-predicted gene pairs. The combined utilization of SLWise and SLAD-CE offers an approach for predicting and analyzing SL interactions in specific cellular contexts. Our findings highlight the potential of SLWise and SLAD-CE in advancing SL-based therapies by improving prediction accuracy and enhancing mechanistic understanding, ultimately contributing to the development of effective precision treatments for cancer.

## Introduction

Synthetic lethality (SL) occurs when non-essential genes become lethal upon simultaneous disruption[1]. This property can be exploited for therapeutic purposes, as targeting SL interactions can selectively kill cancer cells bearing multiple genetic alterations [2-11]. PARP inhibitors, such as olaparib, niraparib, rucaparib, and talazoparib, have been approved for various cancers based on the SL interaction between PARP and BRCA, providing significant benefits to patients in clinics[12-17]. The success of these inhibitors provides sufficient encouragement for the therapeutic application of the SL interactions. Besides, inhibitors of ATR, WEE1, CHK1, and mTOR, the SL partners of tumor suppressor gene TP53, have shown efficacy in clinical development [1, 18, 19]. Inhibitors of PRMT5 and MAT2A, the SL partners of MTAP[8] are also in the clinical stage (https://clinicaltrials.gov/).

Identifying SL interactions is a complex and challenging task. The traditional approaches of genetic screens are expensive, laborious, and time-consuming, limiting the number of genes tested at a time [20]. By combining genome-wide single gene perturbation techniques like RNAi or CRISPR with the inherent abnormalities present in cancer cells, it is possible to uncover certain SL pairs through experimental methods and bioinformatics analyses[21]. However, selective inhibitors targeting specific genes often only benefit a subset of patients with target-labeled markers, indicating that SL interactions likely involve not just two genes but a collective outcome of multiple background genes and target genes. Combination knockout or interference has shown promise in addressing this situation and been gradually implemented[22-32]. However, due to the vast library size, gene pairs need to be filtered based on various criteria, resulting in only a limited number of gene pairs being tested. In response to these challenges, computational approaches that utilize machine learning algorithms and large-scale omics datasets have emerged as promising alternatives for the discovery of SL targets in genome wise scale[33, 34].

Review by Wang et al. provides specific references to machine learning model approaches, databases, and tools for summarization[34]. However, current state of the art ML and DL models in predicting SL interactions have significant limitations.

Firstly, most models integrate the SL gene pairs from the population level as positive labels[35-48]. Population-level data is derived from a set of cell lines or a collection of tumor samples obtained from multiple patients[49]. This approach can lead to reduced reliability due to the phenomenon of cellular context specificity, where gene knockouts may have lethal effects in one cell type but promote growth in another [33, 50-52]. This context specificity has been confirmed through multiple Crispr-Cas9 screens [24, 28]. For instance, Shen et al. (2017) found that only about 10% of SL interactions were shared among three examined cell lines [24]. Ito et al. (2021) reported that only 1.73% of digenic paralog pairs were SL gene pairs in more than 7 of 11 cell lines [28]. In such situations, even when incorporating context-specific biological features such as mutations and gene expression into models, it still becomes challenging to accurately capture the relationship between specific cells/tissues and the predicted SL gene pairs.

In addition, neglecting cell context-specificity undermines the model’s ability to explain predictions by overlooking valuable insights into specific biological processes within cells. To address this, it is vital to incorporate cell context-specific characterization using diverse omics data of the given cell line, including mutation profiles, gene expression patterns, and functional divergence. These data ensure the model capture the context specificity inherent to specific cell lines[5, 53]. Besides, the L1000 Connectivity Map [54] provides crucial information on expression changes of representative genes in response to gene perturbations specific to a given cell line. This resource serves as an excellent context representation to comprehensively characterize the global dynamic network. The cell line-specific model EXP2SL via incorporating L1000 data as its major feature has shown a good performance[55]. However, it also exhibits significant limitations. In contrast to the vast number of approximately 20,000 coding genes, EXP2SL is limited to a few hundred input genes, which hampers its ability to characterize cells and predict genome-wide SL interactions. Additionally, the model’s inability to predict unknown cell lines limits its potential to effectively address the diverse and unknown cells present within the heterogeneity of tumor tissues. Another cell line-specific model, MVGCN-iSL[49], adopts a strategy of incorporating cell line-specific SL gene pairs along with certain cell-independent network features as input. This approach holds promise; however, it still faces challenges when it comes to explaining candidate SL pairs and predicting outcomes in unknown cell lines.

Furthermore, regardless of the various input features considered, translational research and clinical application of these SL pairs face serious challenges. SL models may exhibit high prediction scores, but their difficulty of interpretation and high false positive rates create challenges for researchers and potentially hinder their practical applicability.

To overcome these limitations and challenges, we presented SLWise, a context-specific prediction model for SL interaction. SLWise incorporated cell context-specific features such as gene expression, gene essentiality, and L1000 data, as well as key general features including mutual exclusivity and paralogs known to be important in SL interactions[21, 56, 57]. To gain deeper insights into the underlying mechanisms of predicted SL gene pairs, we propose a novel analysis scheme called SLAD-CE. This approach allows for the identification of essential gene abnormalities and utilizes GSEA to locate cell damage. By incorporating a ground truth evaluation standard and leveraging SLAD-CE, we aim to bridge the gap between deep learning models and practical applications in the field of SL in precision medicine.

## Methods

### Model input feature selection

We select some proven SL mechanisms and incorporated them to our models (Figure 1). Specifically, paralogs, which arise from duplicated sequences of a shared ancestor and often perform similar functions[58], exhibit functional redundancy. Their loss is more common in tumors[59], making them potential precision targets for cancer treatment and an essential dataset for SL discovery[21, 57]. Mutually exclusive mutation patterns suggest incompatible driver mutations in tumorigenesis, indicating a potential source of SL interactions [56]. Identifying mutually exclusive gene pairs from tumor genomes is advantageous for discovering SL interactions[28, 56] and can be utilized in prediction models[37, 60, 61]. In addition, the combination of high-throughput CRISPR-Cas9 screens with the background gene’s low expression (which might be caused by mutation, copy number variation, or epigenetic modification) is also a major logic for SL pair discovery[5, 53, 62]. Furthermore, to understand specific SL interactions in the cellular context, it’s important to consider the dynamic relationship of genes. The L1000 Connectivity Map[54] is a valuable resource that tracks cellular responses and helps characterize specific gene relationships based on expression changes caused by gene perturbations, providing a comprehensive understanding of the cellular context for SL interactions.

**Figure 1.**
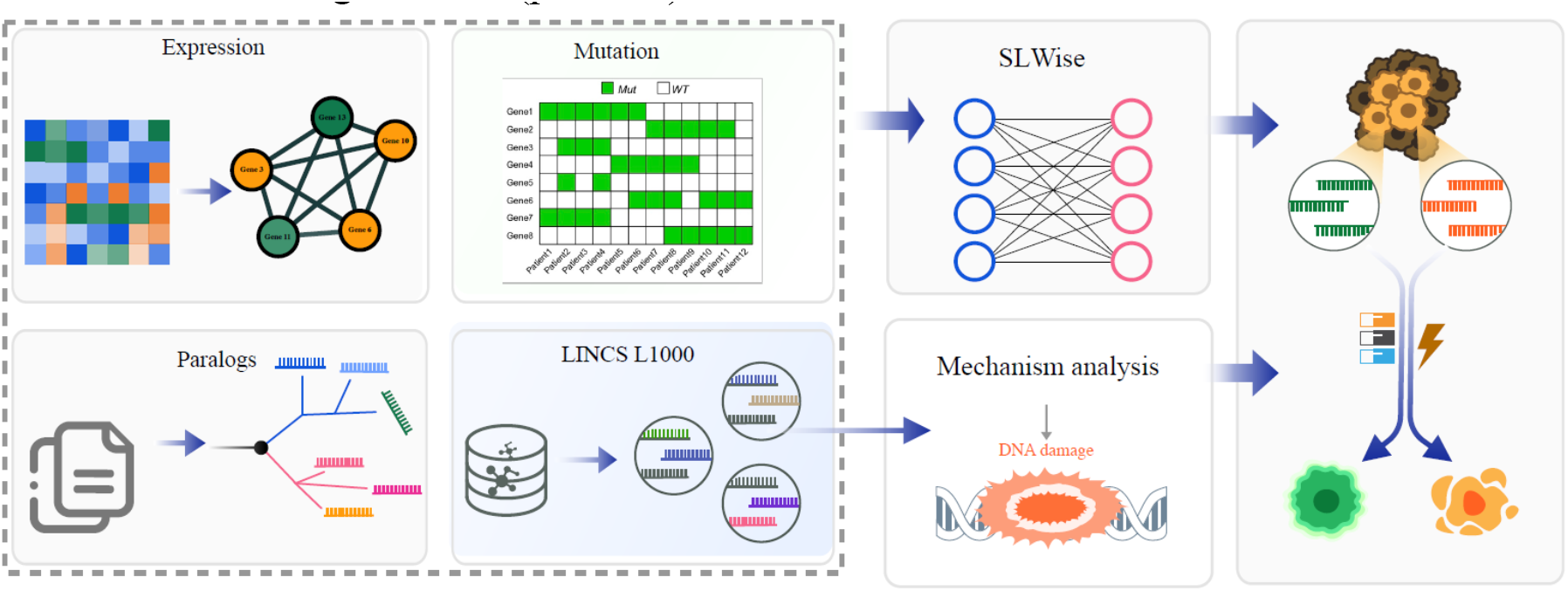
Overview of the cell context-specific SL pair prediction and analysis. The expression, mutation, paralog, and L1000 data as input features were incorporated to SLWise model for SL gene pairs prediction. Subsequently, the predicted SL pairs are subject to mechanism analysis using the SLAD-CE scheme. The combined approach enables the reliable SL gene pair prediction associated with their cell damage mechanisms.

### Data processing

The L1000 data was obtained from the LINCS L1000 project [54], the level 5 data under the CRISPR gene knock-out technology. A 121,489,145*4-dimensional matrix was created by setting the threshold for the absolute value of the fold change to be greater than or equal to 1.5.

Gene effect scores and gene expression profiles were downloaded from the DepMap portal [63](DepMap Public 22Q2 dataset). Genes with low expression were defined as expression values < 1 and expression z-scores < -1.28. Paralog pairs were downloaded from Table S8 in De Kegel et al.[52]. The positive label was obtained from a dual-knockout screening of human tumor cell lines[23, 25, 26, 28] using the GEMINI approach[64]. The p-value for each gene pair was calculated based on the null distribution generated from non-synergistic pairs, which were selected from genes with no expression in the CCLE expression data[28, 64]. Gene pairs with p-values less than 0.05 were classified as positive SL pairs, while those with sensitive lethality scores less than 0 and among the bottom 50% of sensitive lethality scores were classified as negative SL pairs. The number of SL label data is summarized in Table S1. In addition, since the whole genome-size gene-gene matrix is too large and prone to noise, we focus on the enzymatic genes including transferase, hydrolase, and ligase reference the gene screen library by Ito and their collages[28], which comprises a total of 6467 genes. Based on the candidate gene matrix, we evaluated the performance of SLWise.

### An Overview of SLWise

The SLWise is a novel multiplexed GNN model that utilizes multiple biological data sources to generate graphs representing gene and cell information. These graphs are then used to extract relevant features that can be used to predict SL interactions. The SL graph, which contains information about the interactions between genes and cells, serves as the support training data for the model. We employ GraphSage to extract information from our input data. The multi-head transformer is utilized to integrate multiple features derived from disparate input data sources. Finally, the combined features are fed into a fully connected multi-layer perceptron (MLP) for SL interaction prediction.

### Benchmark

We choose commonly used SL prediction models NSF4SL[65], MGE4SL[66] and a cell-line-specific EXP2SL[55] as the benchmark.

- NSF4SL[65]: use two branches of neural networks to learn SL-related gene representations.
- MGE4SL[66]: combine graph neural network and existing knowledge to predict SL pairs.
- EXP2SL[55]: use L1000 data as the input features to predict cell-line-specific SL pairs.

### Analyses of Cellular Context-specific SL Mechanisms

Literature-curated SL pairs and the top-ranked SL pairs from SLWise model were used to decipher the mechanisms underlying the cellular specificity. Each gene of these SL pairs was used as the candidate perturbation gene. L1000 data include many other gene expression data in the given cell line used for candidate gene perturbation. The significantly perturbed gene sets were identified by setting the log2^FC^ > 1 and adjusted p-value < 0.05 in expression compared to the control from L1000 data. For the candidate gene pair including gene A and gene B (Figure 2B), we filtered out the gene_set A that significantly down regulated by gene A, the gene_set B that significantly down regulated by gene B, and gene_set C, the overlapping portion of the gene_set A and gene_set B, which can be down regulated by either gene A or B. The genes in gene set C that both classified as tumor driver genes[67] and having a CERES score less than -0.5, were retained as the essential genes. Then, to localize potential cellular damage, we performed GSEA using clusterProfiler (version 4.4.4) [68] on the subset of genes in gene set C with CERES scores less than -0.5. The enrichment items that achieved statistical significance (p < 0.01) were taken into consideration.

**Figure 2.**
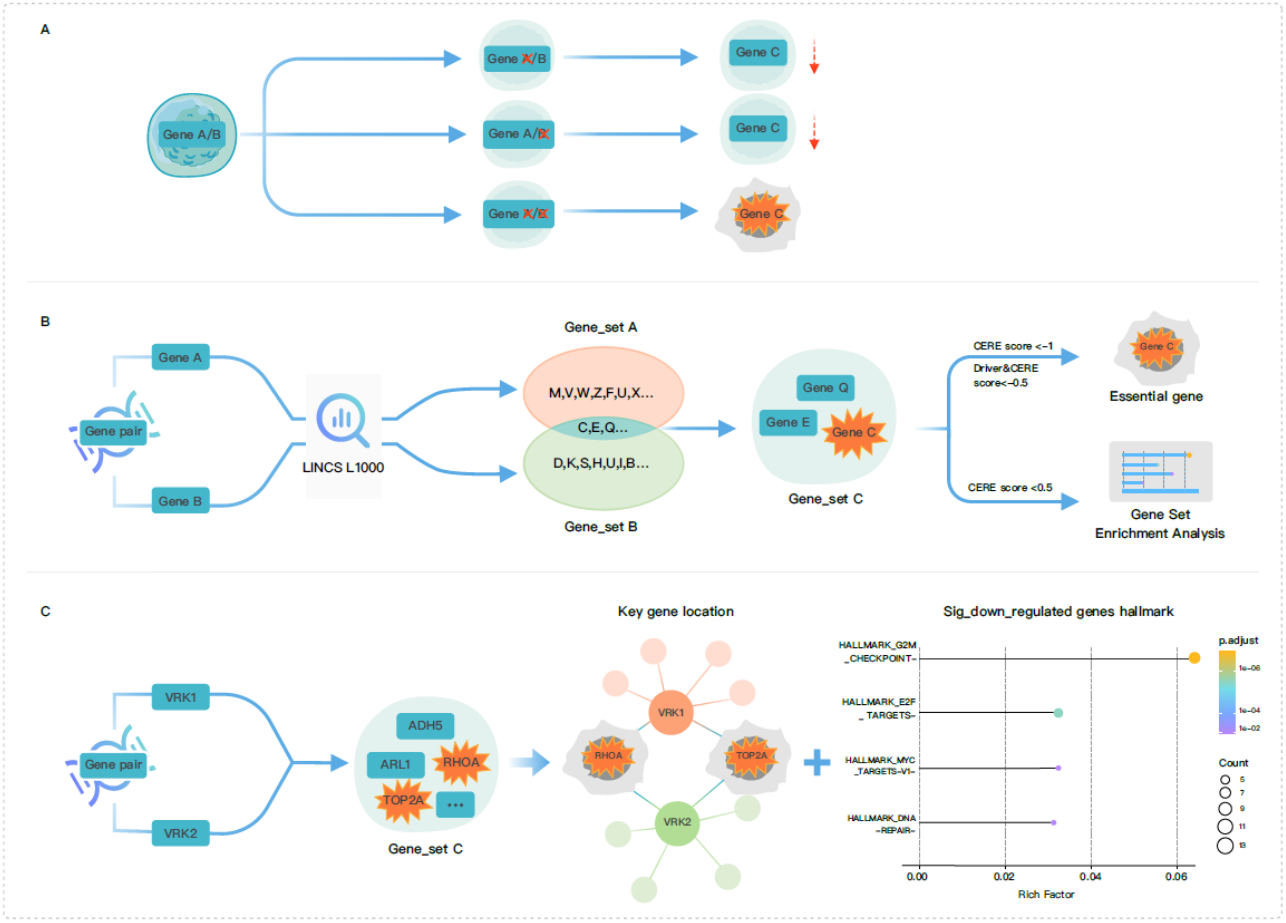
Scheme and a presentative case of mechanism analysis. (A). Mechanism diagram of SL gene pairs induced cell damage. (B). Flowchart depicting the process of identifying essential genes abnormality and assessing damage from candidate SL gene pairs in specific cellular contexts using SLAD-CE. (C). A representative case (VRK1-VRK2) of mechanism analysis process using SLAD-CE.

## Results

### Performance Evaluation of SLWise and other SL Prediction Models

Machine learning models can identify many potential SL interactions with high AUC or F1 scores. However, assessing the practical value of these candidate SL gene pairs remains a perplexing dilemma for researchers. The substantial cost and time required for extensive validation efforts also pose significant challenges in determining the feasibility and practicality of these predicted SL pairs. As a result, a notable gap exists between the models and their actual implementation. To ensure that the prediction results are applicable, it is crucial to evaluate the model’s performance on their a handful of top-ranked candidate genes. Thus, we utilize a ground truth standard to assess the performance of our model by comparing the top-ranked genes (the top 100 and 200) with the actual number of true positive SL gene pairs.

It is clear that SLWise model outperformed the EXP2SL, NSF4SL, and MGE4SL models in accuracy using true positive labels (Table 1). Among the top 100 predictions, SLWise successfully identified 22 true SL pairs in A375 and 2 true SL pairs in HT29 (Table 1, Table S2), while EXP2SL failed to identify any SL pair. Notably, 20 out of 22 true SL pairs in A375 and 1 out of 2 in HT29 exhibited cell context-specificity, meaning that the SL gene pairs had lethal effects in one cell type but not in another due to their different cellular context. Expanding the evaluation to the top 200 predictions, EXP2SL exhibited even lower performance, failing to identify additional SL pairs. In contrast, SLWise identified 30 true SL pairs (27 of which are cell context-specific) in A375, 6 true SL pairs (5 of which are cell context-specific) in HT29, and 0 true SL pairs in A549. Neither NSF4SL nor MGE4SL identified any SL pairs within the top 200 results. Moreover, when extending the analysis to the top 1000 predicted SL pairs, SLWise consistently demonstrated better performance. It successfully identified 30 true SL pairs (27 of which were cell context-specific) in the A375 cell line, 7 true SL pairs (6 of which are cell context-specific) in the HT29 cell line, and 5 true SL pairs (all of which are cell context-specific) in the A549 cell line (Table S3). In contrast, EXP2SL only identified 7 true SL pairs (5 of which are cell context-specific) in the A549 cell line and failed to identify any SL pairs in other cells. MGE4SL identified only 4 true SL pairs, and NSF4SL failed to identify any true pair. Furthermore, we conducted a performance comparison by employing the same approach as the EXP2SL method, focusing on using the same cell line for training and testing. In this evaluation, SLWise still demonstrated better performance over EXP2SL (Table S4). These findings demonstrate the evident superiority of our SLWise model’s accuracy in predicting cell context-specific SL interactions compared to the baseline methods.

**Table 1.** Top100 and 200 predicted SL pairs performance evaluation.

|  |  | Taining SL data | Testing SL data | Labeled SL pairs | Cell context-specific | General |
| --- | --- | --- | --- | --- | --- | --- |
| Top100 | SLWise | A549 | A375 | **22** | **20** | 2 |
|  |  | A549 | HT29 | **2** | **1** | 1 |
|  |  | A375 | A549 | 0 | 0 | 0 |
|  | EXP2SL | A549 | A375 | 0 | 0 | 0 |
|  |  | A549 | HT29 | 0 | 0 | 0 |
|  |  | A375 | A549 | **5** | **4** | 1 |
| Top200 | SLWise | A549 | A375 | **30** | **27** | 3 |
|  |  | A549 | HT29 | **6** | **5** | 1 |
|  |  | A375 | A549 | 0 | 0 | 0 |
|  | EXP2SL | A549 | A375 | 0 | 0 | 0 |
|  |  | A549 | HT29 | 0 | 0 | 0 |
|  |  | A375 | A549 | **5** | **4** | 1 |
|  | NSF4SL | SynLethDB 2.0 |  | 0 | 0 | 0 |
|  | MGE4SL | SynLethDB |  | 0 | 0 | 0 |

For a more intuitive visualization, we have listed several SL gene pairs labeled as positive among the top 100 prediction results (Table 2). It is worth noting that many of them have been validated using low throughput experiments. For example, the paralog pairs such as BCL2L1-MCL1[25, 69] and PARP1-PARP2[70] have been verified for SL interactions. Besides, the combination of BCL2L1 or BCL2L2 knockdown and PARP inhibitor has demonstrated a significant reduction in the viability of certain tumor cells [71]. The combined inhibition of WEE1 and PARP1 has been shown to induce SL interactions, particularly when combined with radiation [72-74]. The co-inhibition of AKT3 and WEE1 has been proven to decrease the development of melanoma[71].

**Table 2.** SL pairs with positive label in the top 100 predicted results.

| Cell line | Gene A | Gene B | Cell context-specific |
| --- | --- | --- | --- |
| A375 | BCL2L2 | UBC | Yes |
|  | BCL2L2 | WEE1 | Yes |
|  | MAPK3 | UBC | Yes |
|  | PARP1 | UBC | Yes |
|  | BCL2L1 | BCL2L2 | Yes |
|  | BCL2L1 | WEE1 | Yes |
|  | BCL2L2 | MAPK3 | Yes |
|  | BCL2L2 | PARP1 | Yes |
|  | MAPK3 | WEE1 |  |
|  | PARP1 | WEE1 | Yes |
|  | MAPK3 | PARP2 | Yes |
|  | PARP1 | PARP2 | Yes |
|  | BCL2L2 | MCL1 | Yes |
|  | MCL1 | WEE1 |  |
|  | AKT3 | UBC | Yes |
|  | AKT3 | BCL2L2 | Yes |
|  | AKT3 | WEE1 | Yes |
|  | AKT3 | PARP2 | Yes |
|  | BCL2L1 | MAPK3 | Yes |
|  | BCL2L1 | PARP1 | Yes |
|  | MAPK3 | PARP1 | Yes |
|  | BCL2L1 | MCL1 | Yes |
| HT29 | MAPK3 | WEE1 |  |
|  | MAPK1 | MAPK3 | Yes |

### Strategy of SL Gene Detection and Cell Damage Evaluation Reveals Cell Context-specific Mechanisms of SL Interactions

We proposed a mechanism analysis strategy, utilizing the logic of synthetic lethal associated gene detection and cell damage evaluation (SLAD-CE), to identify abnormalities in essential genes involved in cell survival and death resulting from the knockout of candidate SL genes. SLAD-CE aims to establish associations between candidate SL gene pairs and cellular damage, providing insights into the underlying mechanisms of SL interactions a cellular context-specific manner. It has been demonstrated that the SL interaction between gene pairs can lead to cell death by disrupting another crucial gene or indicate cellular vulnerability through an additional pivotal gene. For instance, the simultaneous loss of CREBBP and EP300 can trigger the disruption of the MYC gene, ultimately resulting in cell death [75]. Deletion of both NXT1 and NXT2 leads to the rapid degradation of the essential protein NXF1, leading to cell death as well [7]. Through the downregulation of RPS6, the dual inhibition of EGFR and MET induces triple-negative breast cancer cell death[76]. Notably, these genes, including MYC, NXF1, and RPS6, all are essential genes whose disruption or removal can have serious consequences for cells, such as cell death or impaired viability. The essentiality of these genes can be obtained by querying the DepMap website(https://depmap.org/portal/). Based on the aforementioned observations, it becomes evident that discovering the underlying mechanisms of combined effects requires the consideration of at least three genes and the assessment of the essentiality of other genes beyond the first two.

To explore relationships between genes in the context of dual gene knockout, it is imperative to analyze transcriptome data from cells experiencing the corresponding type of perturbation. Currently, such data is scarce. To overcome this limitation, SLAD-CE infers such data from transcriptomes of both single gene knockouts to deduce a three-gene relationship involving gene A, B, and their commonly affected gene C (Figure 2A). To further refine this kind of SL mechanism, we applied a filtering criterion to gene_set C, specifically focusing on essential genes that are driver genes with a CERES score less than -0.5(Figure 2B). This process enables us to build a relationship in which the simultaneous knockout of gene A and B leads to cellular abnormalities, primarily attributed to the dysregulation of essential genes, or serves as an indicator of cellular damage mediated by these genes (Figure 2A).

In our analysis, we investigated a set of 90 validated SL pairs (Table S5) reported in the literature using the A375, A549, and HT29 cell lines. Among these pairs, 46 of these pairs were associated with an essential gene that serves as both a tumor driver and exhibits CERES scores below -0.5. For example, we discovered that knockout of CREBBP and EP300 significantly down regulated six common genes in HT29. Among these genes, only MYC is a driver with a CERES score of -2.07. This finding aligns with the established mechanism by which CREBBP and EP300 induce SL through the abrogation of MYC. Similarly, DUSP4 and DUSP6 were found to significantly down regulate BRAF (with a CERES score of -0.70) in A375 cells harboring the activated BRAF mutation, highlighting their potential role in SL interaction. By employing this approach, we gain valuable insights into how cellular context-specific SL interactions occur.

Furthermore, applying GSEA to genes within gene set C that have CERE scores below -0.5, SLAD-CE is capable to assess cell damage caused by the candidate gene pair. For instance, when examining the relationship between VRK1 and VRK2, we observed that they are associated with the essential gene TOP2A (Figure 2C), a tumor driver gene essential for cell division [77], suggesting a potential SL interaction between VRK1 and VRK2 through the dysfunction of TOP2A. The GSEA analysis of their gene_set C revealed abnormalities in mitotic anaphase and cell cycle checkpoints (Figure 2C), providing additional evidence of the damage caused by this gene pair. This result aligns with the known mechanism wherein the loss of the gene pair leads to G2/M arrest and DNA damage[78]. Moreover, several other gene pairs, such as HDAC1_HDAC2, PARP1_BRCA1, and UBB_UBC, were found to be enriched in G2M_CHECKPOINT, MYC_TARGETS_V1 and DNA_REPAIR (Figure S1). T h e g e n e p a i r M E 2 a n d M E 3 e x h i b i t e d e n r i c h m e n t i n HALLMARK_OXIDATIVE_PHOSPHORYLATION (Figure S1). Meanwhile, all of these gene pairs were associated with a significant down regulation of essential driver genes.

Due to the limitation of data available in cell lines and the specific SL mechanisms considered, this study focused on analyzing three cell lines to determine cell context-specific SL interactions. For instance, regarding VRK1-VRK2, our findings demonstrate that both essential gene abnormalities and significant Hallmark enrichment in cell damage were exclusively observed in the A375 cell line. Similarly, ME2-ME3 and UBB-UBC exhibited a comparable pattern in the A549 and HT29 cell lines, respectively. These results indicated the cell context-specificity of these SL pairs is within the scope of our study. Conversely, the knockout of PARP1 and BRCA1 was associated with essential gene abnormalities, such as RHOA and SUZ12 in the A375 cell line, CDK4 in the A549 cell line, and GMPS in the HT29 cell line. Furthermore, their Hallmark enrichment analysis revealed cell damage occurring in t h e H A L L M A R K _ D N A _ R E P A I R p a t h w a y f o r A 3 7 5 a n d t h e HALLMARK_MYC_TARGETS_V1 pathway for A549. These observations suggest that the gene pair of PARP1 and BRCA1 does not exhibit cell context-specificity within the scope of our study.

In conclusion, the SLAD-CE scheme facilitates the understanding of the cell context-specific SL mechanisms and cellular damages caused by gene pair knockouts. It provides insights into precise SL mechanisms across diverse cell line contexts. Therefore, it could guide the interpretation of the potential new SL interaction identified by our SLWise model.

### SLAD-CE Analysis of the SL Mechanism of gene pairs predicted by SLWise Model

Having validated our mechanism analysis approaches using known SL pairs, we applied SLAD-CE to SL pairs from our SLWise model. In the A375 cell line, we predicted an SL interaction between BCL2L2 and WEE1(Figure 3A). Applying the SLAD-CE approach, we discovered that the knockdown of BCL2L2 and WEE1 in the A375 cell line resulted in abnormalities in two significant driver genes, CNOT9 and RHOA (Figure 3B). It is noteworthy that RHOA plays a crucial role in the growth, progression, and metastasis of various cancer types and has been considered a therapeutic target [79]. Moreover, the cell damage is enriched in mitotic anaphase and M phase, as observed in the Reactome analysis (Figure 3C), and in E2F_targets and G2M checkpoint, observed in the Hallmark analysis (Figure 3D). These findings are consistent with previous reports indicating that the inhibition of WEE1 in tumor cells increases the dependency on BCL2L2[80], providing a plausible explanation for the observed cellular damage. In contrast, in the HT29 cell line, the knockout of BCL2L2 and WEE1 did not result in abnormalities in any essential driver gene, and there was no significant enrichment observed in the Hallmark and Reactome pathways. Additionally, although a significant down regulation of the essential driver gene was detected in the A549 cell line, no significant enrichment was observed in the Hallmark and Reactome pathways.

**Figure 3.**
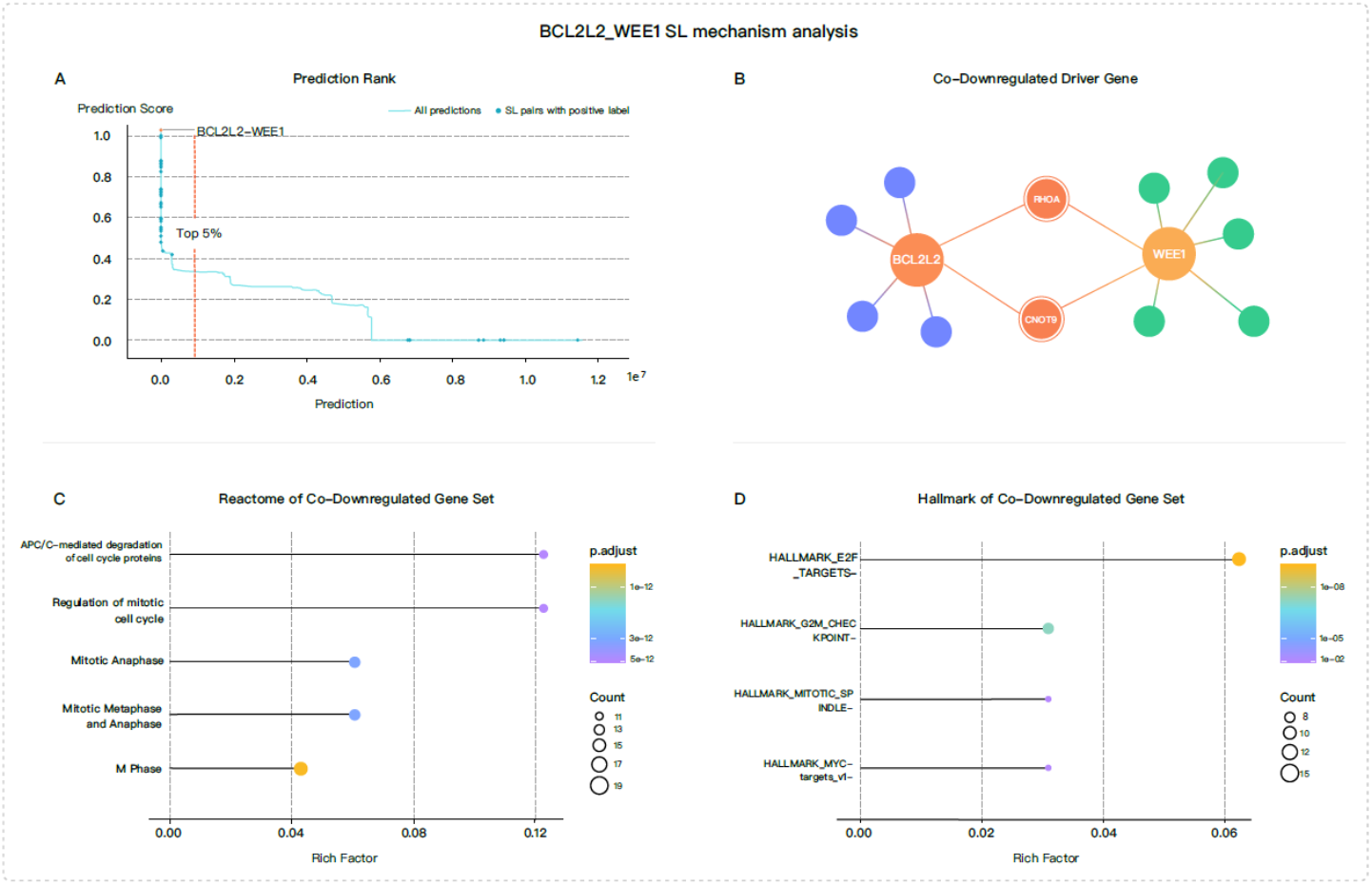
Representative predicted gene pair (BCL2L2-WEE1) and its SL mechanism analysis using SLAD-CE in A375 cell line. (A). The prediction score and rank of all candidate SL gene pairs. The blue dots are SL pairs with positive label. Predictions falling on the left side of the dotted orange line represent the top 5% ranking. (B). Among the downregulated genes of BCL2L2 and WEE1, RHOA and CNOT9 are the common significant driver gene with a CERES score below -0.5 in A375. (C). The Reactome pathway analysis demonstrates the presence of cellular damage. (D). The Hallmark analysis demonstrates the presence of cellular damage.

Similarly, in the HT29 cell line, we observed an SL interaction between UBC and UBE2L6 (Figure 4A). UBC and UBE2L6 encode Ubiquitin and E2 ubiquitin-conjugating enzymes, respectively, which are critical post-translational modifiers involved in maintaining genome stability. Upon analyzing the effects of UBC and UBE2L6 knockout in the HT29 cell line using SLAD-CE, we found that the only essential gene STIL abnormality (Figure 4B). Notably, inhibition of STIL has been shown to suppress tumor progression, indicating its importance in cancer development [81, 82]. Further analysis using Reactome and Hallmark enrichment revealed that the down regulation of UBC and UBE2L6 led to mitotic damage (Figures 4C and D). This suggests that the disruption of UBC and UBE2L6 may induce cellular damage in the HT29 cell line through the inhibition of STIL, leading to impaired mitotic processes. In contrast, in the A375 cell line, no essential driver genes were identified, and no Hallmark or Reactome enrichments were observed.

**Figure 4.**
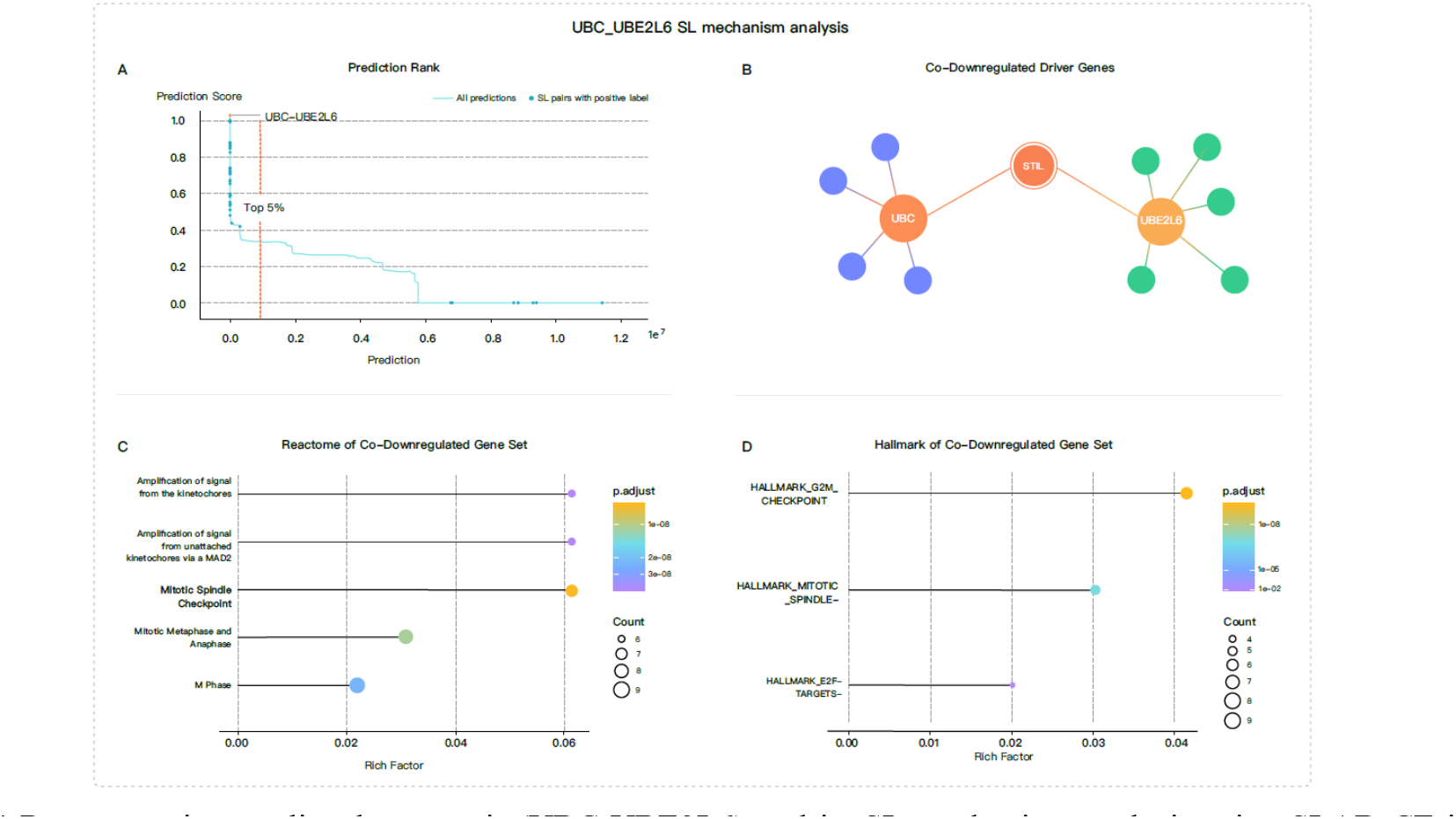
Representative predicted gene pair (UBC-UBE2L6) and its SL mechanism analysis using SLAD-CE in HT29 cell line. (A). The prediction score and rank of all candidate SL gene pairs. The blue dots are SL pairs with positive label. Predictions falling on the left side of the dotted orange line represent the top 5% ranking. (B). Among the downregulated genes of UBC and UBE2L6, STL is the only common significant driver gene with a CERES score below -0.5 in A375. (C). The Reactome pathway analysis demonstrates the presence of cellular damage. (D). The Hallmark analysis demonstrates the presence of cellular damage.

Overall, SLAD-CE analysis highlights BCL2L2-WEE1 and UBC-UBE2L6 pairs as potential SL genes in these cell lines. These findings validate the feasibility and accuracy of the SLWise model in predicting cell context-specific SL interactions.

## Discussion

Applying ML/DL models to real-world applications often faces challenges in interpreting and translating prediction results into application. In this study, we aimed to tackle some of these challenges by incorporaing the SLWise model with the SLAD-CE mechanism analysis scheme, to predict and elucidate cell context-specific SL interactions and their underlying mechanism. To validate model reliability and robustness, we conducted a comparative analysis with baseline models against a ground truth standard. The results demonstrated the superior performance of SLWise against other models in predicting cell context-specific SL interactions. Through detailed case analyses, we unveiled the potential mechanisms underlying newly predicted SL pairs.

Heterogeneity of gene expression levels is a common phenomenon observed in cells with identical DNA, and it can be influenced by various factors such as DNA modification, epigenetic regulation, external stimuli, and cytokines. While the complete understanding of cell context-specific transcription mechanisms remains elusive, it is clear that cell line-specific transcriptional profiles play a crucial role in defining cellular specificity[83, 84]. Transcription data analysis is currently the most widely used method for characterizing and distinguishing cell types[85, 86]. In our model, we leverage gene expression data to investigate the context-specific nature of SL interactions. To capture the dependencies between genes and their potential mechanisms, we proposed a novel scheme called SLAD-CE to locate the essential gene abnormality and cell damage using cell line-specific L1000 and GSEA. This integrated approach allows us to explore plausible mechanisms and evaluate the functional impact of candidate SL pairs in given cells. By analyzing specific examples like BCL2L2-WEE1 in A375 and UBC-UBE2L6 in HT29, we highlight the potential effects and damage caused by these SL pairs in specific cell lines. It is important to acknowledge that the mechanism analyzed using SLAD-CE represents only one among many possible mechanisms, and our SLWise model has the potential to learn beyond what has been discussed here. Notably, although EXP2SL also incorporates the L1000 data, it relies solely on a limited set of genes from L1000 data as its input features, resulting in an incomplete learning of genegene relationships. This limitation could hinder the capture of the fundamental gene interaction logic behind cell line characteristics and SL pairs, making it suboptimal performance for transferability among cell lines.

Obviously, the combination of SLWise model and SLAD-CE scheme provide valuable insights into predicting and analyzing SL interactions. However, they have some limitations that could be addressed through further data expansion and model improvements. Firstly, among the top 200 predicted SL gene pairs for A549, no positive SL labels were observed. This discrepancy may be attributed to the inaccuracy of the model for this specific cell line, possibly derived from the limited availability of positive SL labels (from A375 or HT29, Table S1) during the training process. To overcome this limitation, expanding the dataset with more positive SL labels for various cell lines could improve the performance of the model. Another limitation of SLWise is the lack of transcription and translation regulation information, resulting in a relatively coarse representation of the underlying mechanisms and cellular specificity. To enhance the precision and explanatory power of the model, future versions of SLWise could incorporate gene regulation networks and protein-protein interaction networks to provide a more detailed and comprehensive analysis. Furthermore, considering factors like the nutrient microenvironment and cell-cell interactions will be crucial when applying the model from tumor cells to tumor microenvironment. These additional environmental factors can influence SL interactions and should be considered to obtain a more realistic representation of SL interactions. Additionally, the current version of the model does not integrate mechanisms such as genetic epistasis and regulation by long non-coding RNAs, which could enhance the accuracy of predictions once relevant data becomes available. Cells are highly complex and unique, and the mechanisms underlying SL interactions are diverse and cell context-specific. It is important to acknowledge that the limited mechanistic logic employed in this study cannot capture all possible SL interactions. The focus here is on capturing the damage caused by differential gene expression levels and resulting SL interactions. In future, incorporating more mechanism principles and utilizing them as input features and interpretability tools will help improve the model’s performance and expand its applicability.

This study presents an advancement in the practical application of SL predictive models and provide a analytical scheme to enhance the mechanism interpretation. This progress will ultimately expedite the discovery of SL targets for precision medicine in cancer treatment.

## Acknowledgements

The authors are grateful to all members of the Innovation Center for their helpful discussions. The authors also thank Y. Wang, S. Guo, X. Xiang, J. Zhou, and all members of the StoneWise AI. Inc. for administrative support.

## Author contributions statement

Y.X. and Y.Z. initiated the project. Y.X. designed the SL model input features and mechanism analysis approach, performed the case analysis, and wrote the manuscript. M.P. and K.C. developed the SL model. L.W., G.P., and W.Z. performed the input features processing. Y.X., L.W., and L.Z contributed on data visualization. K.T. assisted in the interpretation of results. All authors provided feedback on the manuscript and approved the submission.

